# Evofosfamide and Gemcitabine Act Synergistically in Pancreatic Cancer Xenografts by Dual Action on Tumor Vasculature and Inhibition of Homologous Recombination DNA Repair

**DOI:** 10.1101/2022.07.04.498761

**Authors:** Yasunori Otowa, Shun Kishimoto, Yu Saida, Kota Yamashita, Kazutoshi Yamamoto, Gadisetti VR Chandramouli, Nallathamby Devasahayam, James B. Mitchell, Murali C. Krishna, Jeffery R. Brender

## Abstract

**Aims:** Pancreatic ductal adenocarcinomas (PDACs) form hypovascular and hypoxic tumors which are difficult to treat with current chemotherapy regimens. Gemcitabine (GEM) is often used as a first line treatment for PDACs, but has issues with chemoresistance and penetration in the interior of the tumor. Evofosfamide, a hypoxia activated prodrug, has been shown to be effective in combination with GEM, although the mechanism of each drug on the other has not been established. We used two mouse xenografts from two cell lines (MIA Paca-2 and SU 86.86) with different tumor microenvironmental characteristics to probe the action of each drug on the other.

**Results:** GEM treatment enhanced survival times in mice with SU.86.86 xenografts (HR =0.35, 95% CI=0.13 to 0.90 p=0.03) but had no effect on MIA Paca-2 mice (HR =0.91, 95% CI=0.37 to 2.25, p=0.84). Conversely, evofosfamide had no effect on SU86.86 mice and did not improve survival times to a statistically significant degree (HR=0.57, 95% CI=0.23 to 1.42, p=0.22). In MIA Paca-2 tumors, which were initially poorly perfused, electron paramagnetic resonance (EPR) imaging showed that oxygenation worsened when treated with GEM, providing a direct mechanism for the activation of evofosfamide by GEM and the effectiveness of evofosfamide and GEM combinations. Sublethal amounts of either treatment enhanced the toxicity of other treatment in vitro in Su86.86 but not in MIAPaca-2. Repair of double stranded DNA lesions was enhanced in the combination treatment in Su86.86 but not MIA Paca-2.

**Innovations:** A possible mechanism for the synergy between evofosfamide and GEM has been proposed.

**Conclusion:** The synergy between GEM and evofosfamide appears to stem from the dual action of GEM’s effect on tumor vasculature and the GEM inhibition of the homologous recombination DNA repair process. The relative importance of each pathway is dependent on the tumor microenvironment and merits further study.

## Introduction

Pancreatic ductal adenocarcinoma (PDAC) is the seventh leading cause of cancer deaths in both men and women [1]. Surgical resection is the only potentially curative treatment; however, 80-90% of patients have a surgically unresectable factor at the time of diagnosis [2]. For patients with an unresectable factor, chemotherapy is a treatment option and gemcitabine (GEM) has been a standard chemotherapy for PDAC since the late nineties [3]. However, while effective initially, PDAC tumors eventually become resistant to GEM within several weeks of treatment [4]. To overcome this resistance problem, combinations of GEM with other cytotoxic and molecular targeted agents have been investigated but have largely failed to show a provable benefit over GEM alone [5-9]. Recently, a combination treatment with leucovorin, 5-FU, oxaliplatin, and irinotecan (FOLFIRINOX) has shown superiority over GEM alone [10]. However, this regime was associated with enhanced toxicity. Therefore, appropriate patient selection is necessary with FOLFIRINOX and GEM monotherapy is still recommended for those whom FOLFIRINOX is contradicted.

Chemoresistance in PDAC tumors stems from a variety of factors, some of which are intrinsic to the tumor and some of which are adaptive responses to treatment. One intrinsic barrier to treatment is the abnormal and leaky vasculature system of PDAC tumors [11] produced as a result of the secretion of antiangiogenic factors [12-15] and compression of the blood vessels by the tumor stroma, which results in poorly perfused chemoresistant regions within the tumor where drug delivery is expected to be inadequate. The hypoxia that results activates the HIF1 transcription factor, which initiates a survival program that upregulates pyrimidine biosynthesis through increased flux through the pentose phosphate pathway. The deoxycytidine triphosphate (dCTP) produced serves as an effective competitive inhibitor for GEM.

A treatment which either targets the poorly perfused regions where GEM cannot penetrate easily or which modifies the tumor microenvironment to decrease the activation of HIF1 may act synergistically with GEM by eliminating potential areas of regrowth or decreasing the competitive inhibition of GEM. The surviving hypoxic regions are of particular concern as hypoxia is correlated with rapid tumor growth in PDAC tumors and a poorer prognosis.[16] The investigational drug Evofosfamide is of special interest as it specifically targets hypoxic areas.[17] Evofosfamide is administered as a prodrug which is reduced *in vivo* to form an unstable nitro anion radical. The fate of the radical is dependent on the local oxygen concentration.[18] In highly hypoxic regions (below 15 mm Hg), the nitro anion undergoes unimolecular fragmentation to release a cytotoxic alkylating agent. In the presence of oxygen, this reaction is reversed and the nitro anion radical is restored to its original state and the drug is inactive.

Evofosfamide is unlikely to be effective as a monotherapy as its ability to target normoxic areas of the tumor is limited by the diffusion of the alkylating agent, which rapidly crosslinks with DNA and other macromolecules. The MAESTRO Phase III trial for unresectable pancreatic ductal adenocarcinoma showed a statistically significant increase in the secondary endpoint of progression free survival for evofosfamide / gemcitabine combination therapy over gemcitabine alone (5.5 months vs. 3.7 months, p = 0.004), [19] in line with several preclinical studies that have shown that combination therapies with evofosfamide have a benefit in PDAC.[18, 20-26] However, the primary endpoint of overall survival was not met in this study.

The extent of the benefit from evofosfamide in a combination therapy is likely to depend on the tumor microenvironment, as can be seen in preclinical models with differing sensitivity to GEM and evofosfamide. MIA Paca-2 PDAC xenografts has a low tumor partial oxygen pressure (pO_2_) and respond to evofosfamide while SU.86.86 xenografts, which have a more functioning vascular system and lower oxygen consumption, do not [22, 27, 28]. On the other hand, SU.86.86 xenografts respond to GEM treatment while MIA Paca-2 xenografts does not.[29] Interestingly, both PDAC xenografts respond well to combination treatment with evofosfamide and GEM, [21, 24] but the underlying mechanism remains unknown.

Non-invasive molecular imaging techniques enables to monitor changes in tumor microenvironment. Electron paramagnetic resonance (EPR) imaging is useful to noninvasively and quantitatively obtain in vivo three-dimensional pO_2_ maps and fractional volume of the hypoxic region of the tumor.[30, 31] In addition, conventional MRI using Gd-DTPA and ultra-small superparamagnetic iron oxide (USPIO) will show the change in the intratumor blood perfusion and blood volume in response to treatment.

## Materials and Methods

### In vitro cell viability

Exponentially growing cells were seeded into 96-well plates at a density of 10,000 cells/well 24 h before the addition of evofosfamide and/or GEM. After the drug addition, the cells were incubated under aerobic, air-equilibrated conditions for 48 h at 37°C in a standard tissue culture incubator. Cell cytotoxicity was assessed using CellTiter-Glo Luminescent Cell Viability assay (Promega Corporation, Madison, USA). The luminescent signal was measured using GloMax luminescence detector. Data were presented as proportional viability (%) by comparing the treated group with the untreated cells, the viability of which is assumed to be 100%. Low and high dose of evofosfamide and GEM were determined according to approximately 90% and 50% dose of viable cells, respectively.

### Measurement of oxygen consumption rate (OCR)

The oxygen consumption rate (OCR) was analyzed on a XF96 Extracellular Flux Analyzer (Agilent Technology). Twenty thousand cells were plated into each well of a 96-well plate and cultured overnight. Cells were treated with GEM (0.1, 1 and 1000 µM) and OCR changes were monitored.

### Mice and tumors

Cells were routinely cultured in RPMI 1640 with 10% fetal calf serum. The tumors were formed by injecting 2 × 10^6^ subcutaneously into the right hind legs of female athymic nude mice. Tumor-bearing mice were treated daily with the intraperitoneal administration of 50 mg/kg evofosfamide (Threshold Pharmaceuticals) 5 days a week or GEM (Lilly) 150 mg/kg a week when tumor size reached approximately 400 mm^3^. For combination treatment, GEM was injected 3 hours after the injection of evofosfamide. All imaging experiments were performed at the timing of pre-treatment (before the first injection on day 1) and post-treatment (1 h after the 5^th^ treatment on day 5). In the imaging experiments, mice were anesthetized by isoflurane inhalation (4% for inducing and 1%–2% for maintaining anesthesia) and positioned prone with their tumor-bearing legs placed inside the resonator. During EPRI and MRI measurements, the breathing rate of the mouse was monitored with a pressure transducer (SA, Instruments, Inc.) and maintained at 60 ± 20 breaths/min. Core body temperature was maintained at 36°C ± 1°C with a flow of warm air.

### EPRI

Technical details of the EPR scanner and oxygen image reconstruction were described previously [32]. Parallel coil resonators tuned to 300MHz were used for EPRI. After an animal was placed in the resonator, an oxygen-sensitive paramagnetic trityl radical probe, OX063 (1.125 mmol/kg bolus), was injected intravenously under isoflurane anesthesia. The free induction decay (FID) signals were collected following the radiofrequency excitation pulses (60 ns) with a nested looping of the x, y, and z gradients, and each time point in the FID underwent phase modulation, enabling 3D spatial encoding. Because FIDs last for 1 to 5 ms, it is possible to generate a series of T2* maps, i.e., EPR line width maps, which linearly correlate with the local concentration of oxygen if the concentration of Ox063 is low enough to avoid the contribution of self-line broadening, and to allow pixel-wise estimation of tissue pO_2_. The repetition time was 8.0 µs. The number of averages was 4,000. After EPRI measurement, corresponding anatomic T2-weighted MR images were collected with a 1T scanner (Bruker BioSpin MRI GmbH).

### Blood volume imaging

DCE-MRI studies were performed on a 1 T scanner (Bruker BioSpin MRI GmbH). For blood volume (BV) calculation, spoiled gradient echo sequence images were collected before and 5 minutes after injection of ultra-small superparamagnetic iron oxide (USPIO) contrast (2 μL/g of body weight). The imaging parameters included the following: FOV = 28 × 28 mm; matrix = 128 × 128; echo time (TE) = 3.5 ms; TR = 200 ms; and number of average = 12. The percentage tumor BV was estimated as described previously [33].

### Histological assessment

Tumor tissues were excised, frozen with Tissue-Tek O.C.T. compound (Sakura Finetek USA Inc.) by ultra-cold ethanol and sectioned (10 mm) using a cryostat, with the sections being thaw-mounted on glass slides. Sections were fixed with cold acetone for 10 minutes. The nonspecific-binding sites on sections were then blocked using Protein Block Serum-Free reagent (Dako North America Inc.) for 10 minutes at room temperature. For *in vitro* γH2AX staining, 250,000 cells are seeded on 35 mm dishes 24 hours before treatment. 24 hours after the treatment with evofosfamide and/or gemcitabine at the indicated concentration, the dishes were washed with PBS and fixed with 4 % paraformaldehyde, followed by same staining using anti-γH2AX antibody (Cell Signaling inc.; 1:1000) overnight at 4° C and Alexa Fluor 488 anti-rabbit secondary antibody (Invitrogen; 1:500) for 1 h at room temperature. The stained dishes were scanned using a BZ-9000 microscope (Keyence), and the fraction of immunostain-positive cells was analyzed. The stained slides were scanned using a BZ-9000 microscope (Keyence), and the immunostain-positive area was quantified using ImageJ software (downloaded from https://imagej.nih.gov/ij/).

### Statistical analysis

Data were expressed as the means ± standard error. The significance of the differences between groups were analyzed using the Student *t* test. The log-rank test was used to compare the distribution of Kaplan–Meier survival curves. The event for the Kaplan–Meier test was death or the tumor 2000 mm^3^, mice requiring early or unscheduled euthanasia due to unrelieved pain/distress were treated as censored. *p* < 0.05 was considered statistically significant.

## Results

### Evofosfamide and gemcitabine act synergistically in *vivo*

To see the effect of the combination treatment *in vivo*, we evaluated two pancreatic cancer xenograft models, MIA Paca-2 and SU.86.86. These tumor models are known to respond to either evofosfamide (MIA Paca-2) or GEM (SU.86.86) but not to both treatments individually. The treatment was started when the tumor reached approximately 400 mm^3^ and treated with either or both evofosfamide and GEM by i.p. injection. Evofosfamide was injected 5 days of week and GEM was injected once a week for 4 cycles. For combination treatment, evofosfamide was injected 3 h prior to GEM injection.

As individual treatments, evofosfamide and GEM were weakly selective for specific PDAC cell lines (Fig. 1). GEM monotherapy increased survival times (HR =0.35, 95% CI=0.13 to 0.90 p=0.03) and reduced tumor growth in SU.86.86 xenografts but had no effect on MIA Paca-2 tumors survival times (HR =0.91, 95% CI=0.37 to 2.25, p=0.84) (Fig. 1A). Evofosfamide monotherapy alone did not improve survival times to a statistically significant degree (HR=0.57, 95% CI=0.23 to 1.42, p=0.22) and had no measurable effect on SU.86.86 tumors (HR=1.44, 95% CI=0.56 to 3.80, p=0.45). Combination therapy of evofosfamide and GEM improved survival times in both MIA Paca-2 and SU.86.86 and was significantly more effective than either GEM or evofosfamide in both models (Table S1). To consider possible synergistic effects between GEM and evofosfamide, we built Cox regression models considering the effect of GEM and evofosfamide separately and an interaction term between them (S(t)∼ GEM + Evofosfamide + GEM: Evofosfamide) and compared it to a model considering only the additive effect of both treatments (S(t)∼ GEM + Evofosfamide). In this manner, we could confirm a synergy existed between GEM and evofosfamide in the SU.86.86 model (likelihood ratio of the interacting model over the additive model = 5.16, p=0.03) but not in MIA Paca-2 (likelihood ratio 2.38, p=0.12).

**Figure 1.**
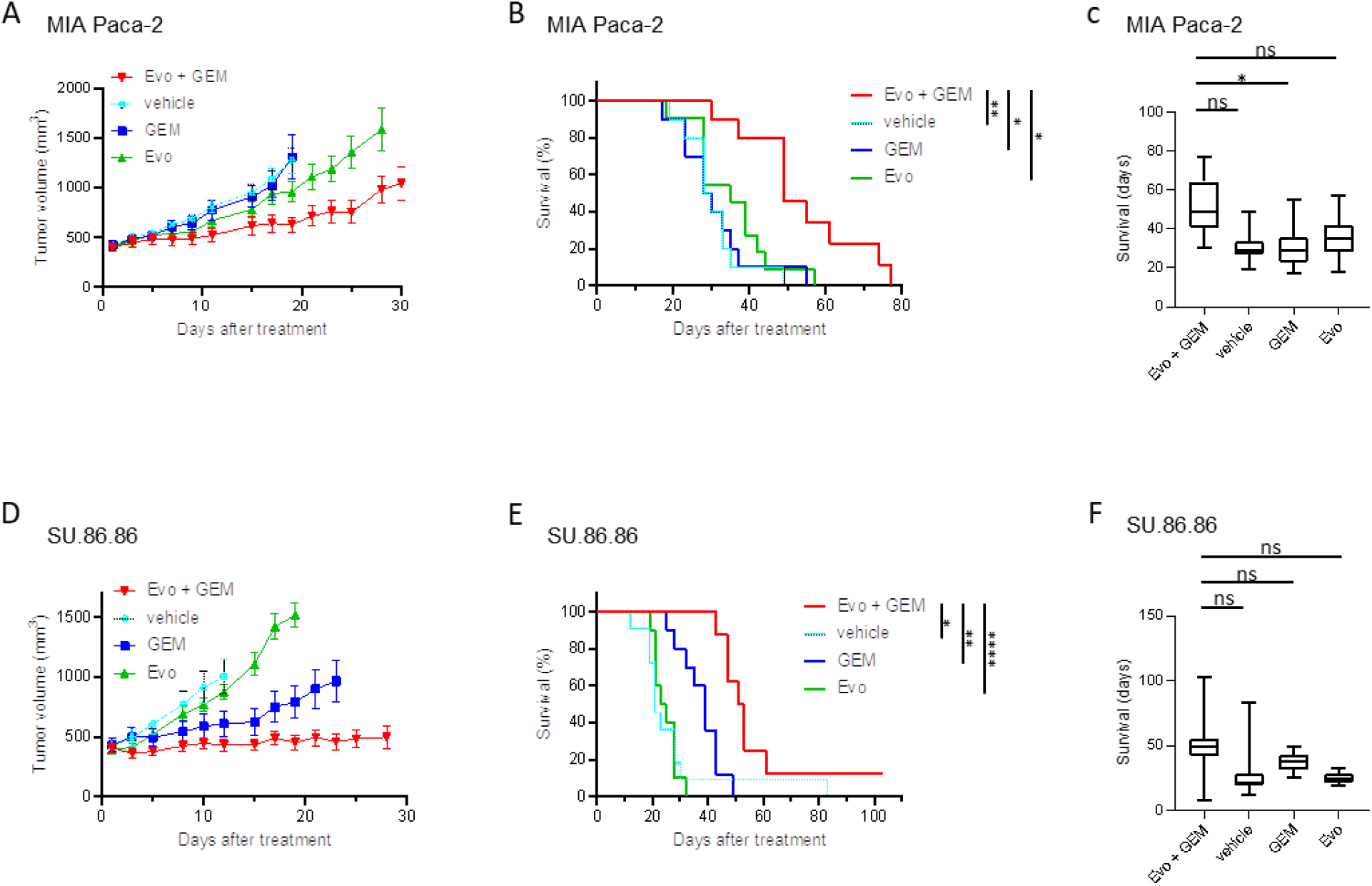
Tumor growth and survival of combination treatment in two murine tumor models with different sensitivity. (**A** and **D**) Growth kinetics of each tumor. MIA Paca-2 or SU.86.86 inoculated mice were treated with either a combination of gemcitabine (GEM) and evofosfamide or GEM or evofosfamide or vehicle (n = 10 per group). Data are shown as mean ± SE at each time point. (**B** and **E**) Kaplan-Meier survival curve for each tumor (MIA Paca-2; n = 10-11 per group, SU.86.86; n =10-11 per group). Survival refers to the time before reaching the maximally allowed tumor volume of 2,000 mm^3^. (**C** and **F**) Bar plot of survival from **B** and **E**. Statistical significance between groups was determined by log-rank test for **B** and **E** and by Tukey-Kramer test for **C** and **F**. \**p* < 0.05; \*\**p* < 0.01; \*\*\*\**p* < 0.0001

### Cell cytotoxicity of evofosfamide and gemcitabine in *vitro*

The differential treatment response between SU86.86 and MIA Paca-2 suggests some critical aspect of pharmacology differs for both GEM and evofosfamide in the two cell lines, which can stem from either a difference in the tumor microenvironment or intracellular differences in metabolism or target sensitivity independent of the tumor stroma. To eliminate the effect of the tumor microenvironment, we tested the cytotoxicity of each tumor model *in vitro* by an ATP based cell viability assay (Fig. 2A). *In vivo*, GEM was effective in the SU.86.86 model but had little to no effect on MIA Paca-2 tumor growth or survival times. *In vitro*, MIA Paca-2 had nanomolar sensitivity to GEM (IC_50_ = 59.2 nM) but was almost two orders of magnitude less effective on SU.86.86 tumor cells (IC_50_ = 3605.9 nM). The strong selectivity GEM shows towards SU86.*86 in vivo* was reversed in *in vitro*, suggesting differences in pharmacodynamics or secreted factors from the stroma that only exist *in vivo* are dominant in determining GEM sensitivity.

**Figure 2.**
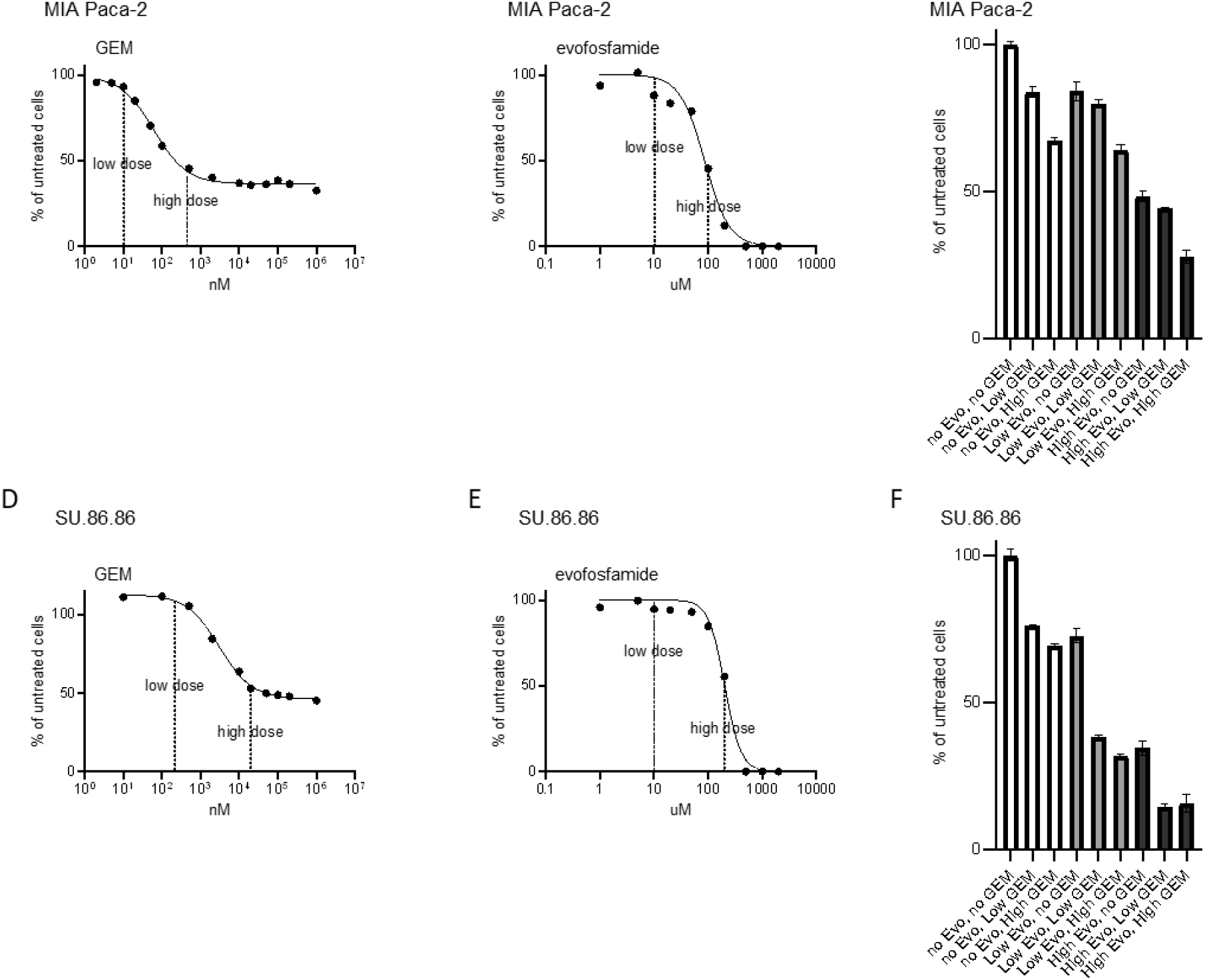
In vitro cytotoxicity evaluated after 48 h of incubation with combination treatment. (**A** and **D**) Dose response curve of MIA Paca-2 and SU.86.86 cells for gemcitabine (GEM) single administration (MIA Paca-2; n = 3 per group, SU.86.86; n = 3 per group). (**B** and **E**) Dose response curve of MIA Paca-2 and SU.86.86 cells for evofosfamide single administration (MIA Paca-2; n = 3 per group, SU.86.86; n = 3 per group). (**C** and **F**) Incubation with combination of GEM and evofosfamide showed a synergistic effect in vitro (n = 3 per group).

Evofosfamide is a hypoxia activated prodrug and is therefore expected to have minimal activity in cell culture under normoxic conditions. Similar to previous reports, we saw some residual micromolar activity of evofosfamide in vitro which was similar in MIA Paca-2 and SU.86.86 tumor cells (IC_50_ =96.3 µM and 216.7 µM, respectively), despite the strong selectivity evofosfamide shows towards MIA Paca-2 *in vivo*. Like GEM, evofosfamide sensitivity *in vivo* is likely dominated by the tumor microenvironment.

A substantial fraction of cells were resistant to GEM in this assay even after treatment for 48 hours at high micromolar concentrations, suggesting a strong chemoresistant component exists in both cell lines that is independent of the tumor microenvironment. A simple explanation for the synergistic effect of evofosfamide and GEM is that evofosfamide kills this resistant fraction, suppressing potential recurrent growth. To explore the possible synergistic effects of GEM and evofosfamide, we tested the ability of low, sublethal concentrations (IC10) of evofosfamide and GEM to sensitize cells to the other treatment.

MIA Paca-2 cells showed no sensitization effect *in vitro*: the addition of sublethal doses of GEM and evofosfamide had no effect on lethal (IC50) of doses of the other treatment, suggesting the synergy present in Fig. 1 may be purely an effect of the tumor microenvironment. Both evofosfamide and GEM treatments, on the other hand, were strongly affected by sublethal concentrations of the other treatment in SU86.86 cells. Sublethal doses of GEM and evofosfamide decreased viability more than the additive effect of either treatment, suggesting the synergy *in vivo* between evofosfamide and GEM is at least partly present *in vitro* for SU86.86.

### Dynamic Changes in tumor oxygen level during tumor progression

Evofosfamide is a hypoxia activated drug whose effectiveness depends on the existence of a hypoxic fraction within the tumor. To evaluate dynamic changes in hypoxia during tumor growth, we acquired baseline EPR oximetry images over a five-day period in the absence of treatment. Representative merged images of anatomy from T_2_-weighted MRI scans and the corresponding EPR-based pO_2_ maps and corresponding frequency histograms of pO_2_ values before and after 5^th^ treatment of vehicle for MIA Paca-2 and SU.86.86 tumors are shown in Figure 3A and 3B and Figure 3E and 3F, respectively. In line with previous reports in the literature, MIAPaca-2 tumors were initially more hypoxic than SU86.86 tumors. However, this difference decreased as SU86.86 tumors uniformly grew more hypoxic while the median pO_2_ and hypoxic fraction of MIA Paca-2 largely remained stable during the five-day vehicle treatment period. In the absence of treatment, the initially normoxic environment of MIA Paca-2 had largely converged to the hypoxic environment of SU86.86.

**Figure 3.**
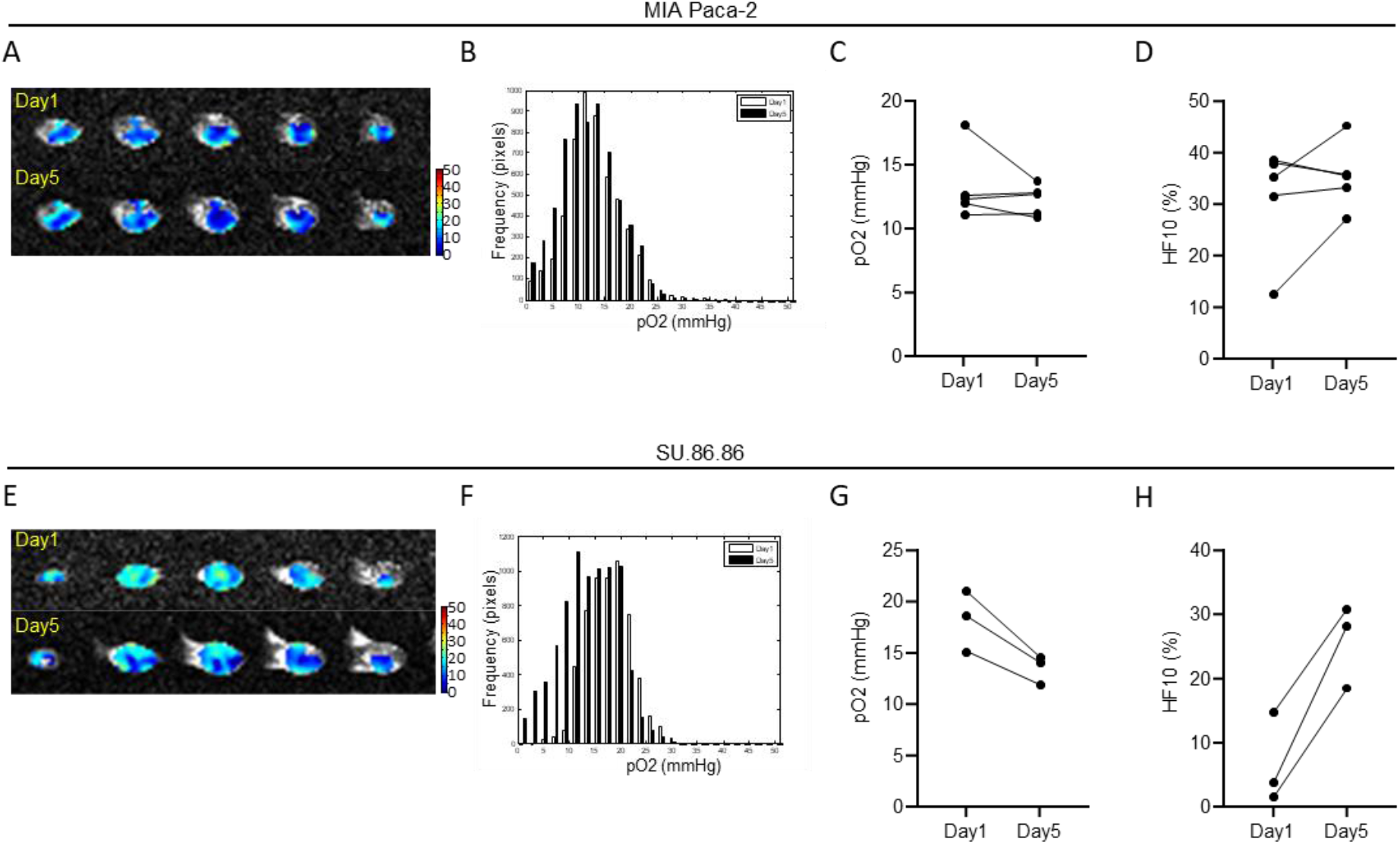
Change in tumor oxygenation using EPR oximetry. (**A**-**H**) EPR imaging in two murine tumor models. Tumor bearing mice treated with vehicle were scanned before and after 5^th^ treatment (MIA Paca-2 vehicle n = 5, SU.86.86 vehicle n = 3). (**A** and **E**) Representative images from MIA Paca-2 and SU.86.86 tumors taken before and after 5^th^ treatment. (**B** and **F**) The frequency histogram of pO_2_ values before and after 5^th^ treatment. (**C** and **G**) The difference of pO2 after treatment with vehicle in MIA Paca-2 and SU.86.86 tumors. (**D** and **H**) The difference of ΔHF10 after treatment with vehicle in MIA Paca-2 and SU.86.86 tumors.

### GEM induces hypoxia leading to a synergistic effect with evofosfamide in MIA Paca-2 tumors

Because the efficacy of evofosfamide is strictly dependent on the hypoxic fraction of the tumor, a decrease in oxygen delivery or an increase in oxygen consumption from GEM treatment would provide a direct mechanism for the synergistic effect of GEM on evofosfamide. Since, unlike SU.86.86, evofosfamide was only effective in MIA Paca-2 in the context of GEM treatment (Fig. 1), we evaluated changes in tumor oxygenation in MIA Paca-2 xenografts after treating with GEM by EPR imaging. EPR-based pO_2_ maps and corresponding frequency histograms of pO_2_ values before and after 5^th^ treatment of GEM are shown in Figure 4A and 4B. To assess the effect of GEM treatment, we calculated the partial oxygen pressure after treatment ΔpO_2_ (ΔpO_2_ = post-treatment pO_2_ − pretreatment pO_2_) and the change of the hypoxic fraction below 10 mmHg ΔHF10 (ΔHF10 = post-treatment HF10 − pretreatment HF10). Median ΔpO_2_ levels decreased compared to the vehicle control, although this trend was not statistically significant (Fig. 4C). The fragmentation of the evofosfamide prodrug is exponentially dependent on oxygen, however, and changes in normoxic or slightly hypoxic cells are unlikely to result in significant releases of the active Br-IPM cross-linking moiety. The deeply hypoxic fraction capable of activating evofosfamide (ΔHF10) significantly increased after GEM treatment (Fig. 4D).

**Figure 4.**
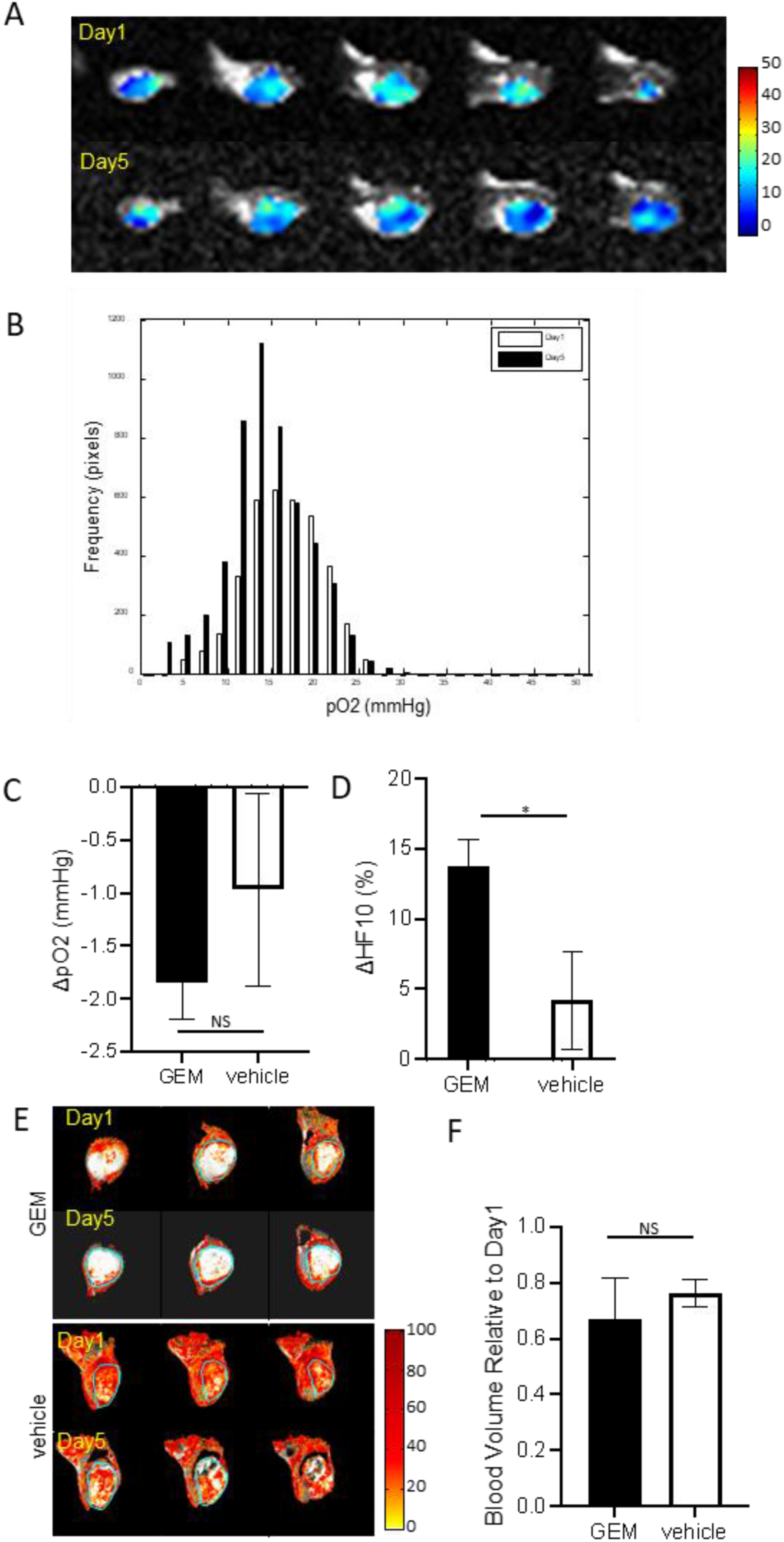
Combination treatment on MIA Paca-2 tumors. (**A** and **B**) Representative EPR images and histogram before and after the 5^th^ treatment of GEM. (**C** and **D**) The difference of ΔpO_2_ (**C**) and ΔHF10 (**D**) between treatment groups (GEM n = 6, vehicle n = 5). (**E**) Representative blood volume images taken before and after 5^th^ treatment (GEM n = 5, vehicle n = 6). (**F**) The blood volume of Day5 relative to Day1. Data are shown as mean ± SE. \**p* < 0.05; NS, nonsignificant.

Hypoxia can result from either an increase in oxygen demand or from a decrease in oxygen supply to the affected areas of the tumor. *In vitro* analysis by Seahorse XF96 Extracellular Flux Analyzer revealed no change in oxygen consumption by MIA-Paca 2 within the first hour after treatment of up to 1 mM GEM and only a transient decrease in the extracellular acidification rate at high concentrations of GEM that resolved within the first 30 minutes (Figure S1). An increase in oxygen demand due to changes in tumor metabolism after GEM treatment is therefore unlikely to be responsible for the increase in the hypoxic fraction, although metabolic contributions from the stroma cannot be ruled out.

To investigate changes in oxygen delivery we employed USPIO imaging, which, when imaged shortly after delivery of the USPIO agent, evaluates changes in blood volume (BV) that can occur through vasoconstriction or changes in patency of the vasculature in response to GEM treatment. It therefore provides a measure of both the overall delivery of oxygen and possible penetration of evofosfamide deep within the tumor. Figure 4I shows BV images calculated from the pre- and post-contrast MRI signal intensity at pre-treatment on day 1 and 1 hour after post 5^th^ treatment on day 5. There was no difference in the summed blood volume within the tumor between GEM and vehicle treated mice (Fig. 4F), indicating a change in blood vessel patency was not responsible for the increase in hypoxia after GEM treatment.

### Evofosfamide treatment is associated with improved oxygenation and blood volume in SU.86.86 tumors

SU86.86 showed the same synergy as MIA PaCa2 but in the opposite direction – in SU86.86 GEM was only effective in the context of evofosfamide treatment. To probe possible changes in the tumor microenvironment in SU86.86 in response to evofosfamide, we first conducted EPR imaging to see changes in tumor oxygenation after treatment. EPR-based pO_2_ maps and corresponding frequency histograms of pO_2_ values before and after 5^th^ treatment of evofosfamide are shown in Figure 5A and B. While evofosfamide did not suppress tumor growth or mouse survival, evofosfamide arrested the decrease in oxygen levels (Fig. 5C) and the increase in the hypoxic fraction that occurred in the control during the treatment period and (Fig. 5D). This change is associated with also associated with a positive change in blood volume. Similar to oxygen levels, blood volume decreased strongly in the control during the treatment period but remained nearly constant throughout the treatment in the Evofosfamide sample (Fig. 5E and 5F).

**Figure 5.**
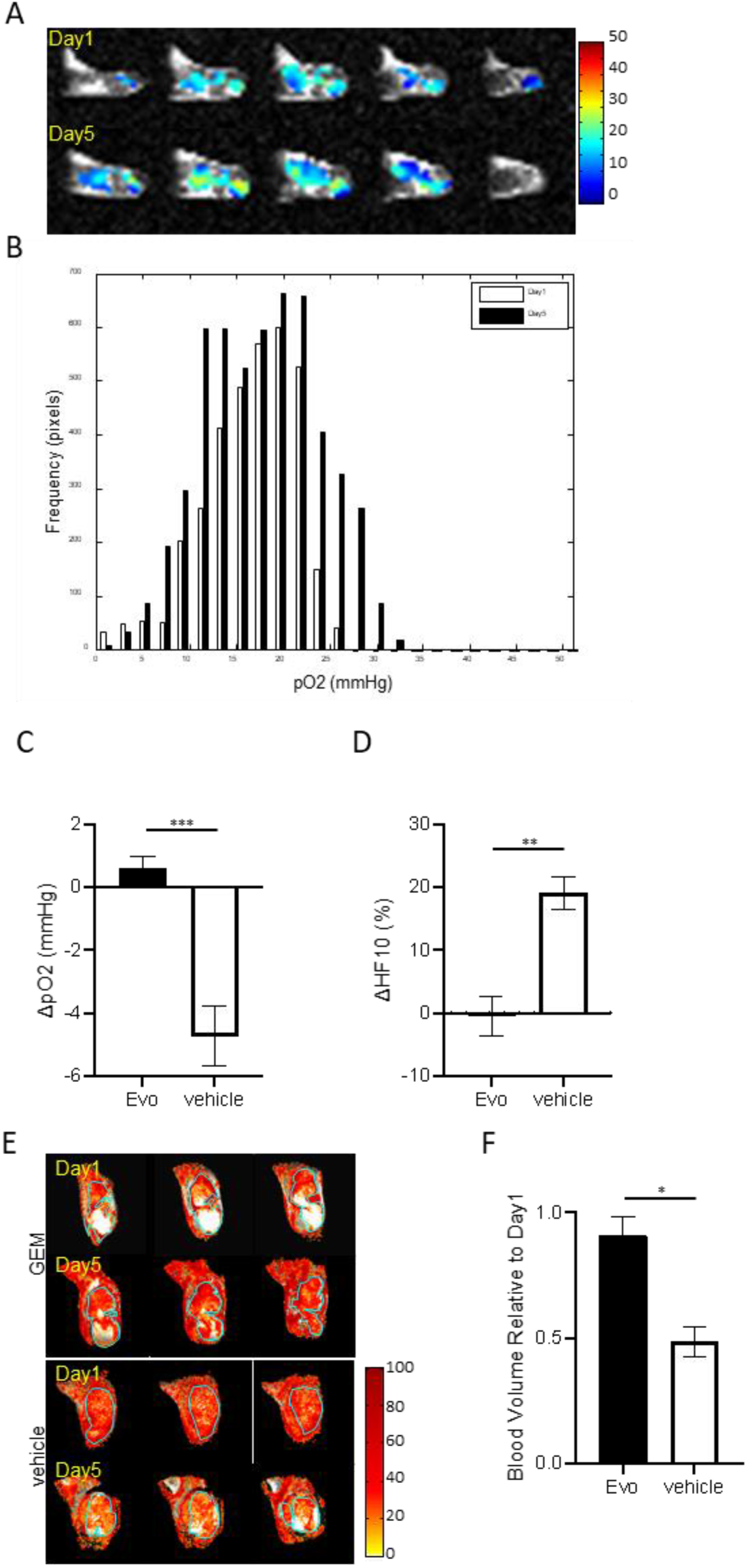
Combination treatment on SU.86.86 tumors. (**A** and **B**) Representative EPR images and histogram before and after the 5^th^ treatment of evofosfamide. (**C** and **D**) The difference of ΔpO2 (**C**) and ΔHF10 (**D**) between treatment groups (evofosfamide n = 5, vehicle n = 3). (**E**) Representative blood volume images taken before and after 5^th^ treatment (evofosfamide n = 7, vehicle n = 4). (**F**) The blood volume of Day 5 relative to Day 1. Data are shown as mean ± SE. Statistical significance between groups was determined by Student *t* test. \**p* < 0.05; * * *p* < 0.01; * * * *p* < 0.001; NS, nonsignificant.

Unlike MIA-PaCa-2, the synergy between GEM and evofosfamide in SU.86.86 was at least partly retained *in vitro* (Fig. 2F), which suggests intracellular biochemistry is largely responsible for the synergy. Evofosfamide causes extensive double stranded DNA lesions with most of that damaged repaired through homologous recombination, [28] a process which is known to be inhibited by GEM.[34-36] We used a histological marker for double stranded breaks, phosphorylated histone γ-H2AX,[37] to measure the number of unrepaired double stranded breaks that occur in adherent SU.86.86 cells after a sublethal dose of GEM, evofosfamide or a combination of the two. (Fig. 6) Neither GEM or evofosfamide showed a strong capacity to induce double stranded breaks *in vitro* in SU.86.86 when used as a monotherapy at these concentrations (0.23 ± 0.07 for GEM at 2000 nM and 0.20 ± 0.07 for evofosfamide at 100 μM). When used at the same concentrations as the monotherapies, the combination of the two drugs was significantly more effective than either drug alone (p= 0.0002 for GEM and 0.0003 for evofosfamide) or the sum of both drugs acting independently. In MIA Paca-2, which did not retain the *in vivo* synergy between GEM and evofosfamide *in vitro*, did not show the same synergy (Fig. S3) implicating homologous recombination as a likely mechanism for GEM and evofosfamide synergy in SU86.86 but not MIA Paca-2.

**Figure 6.**
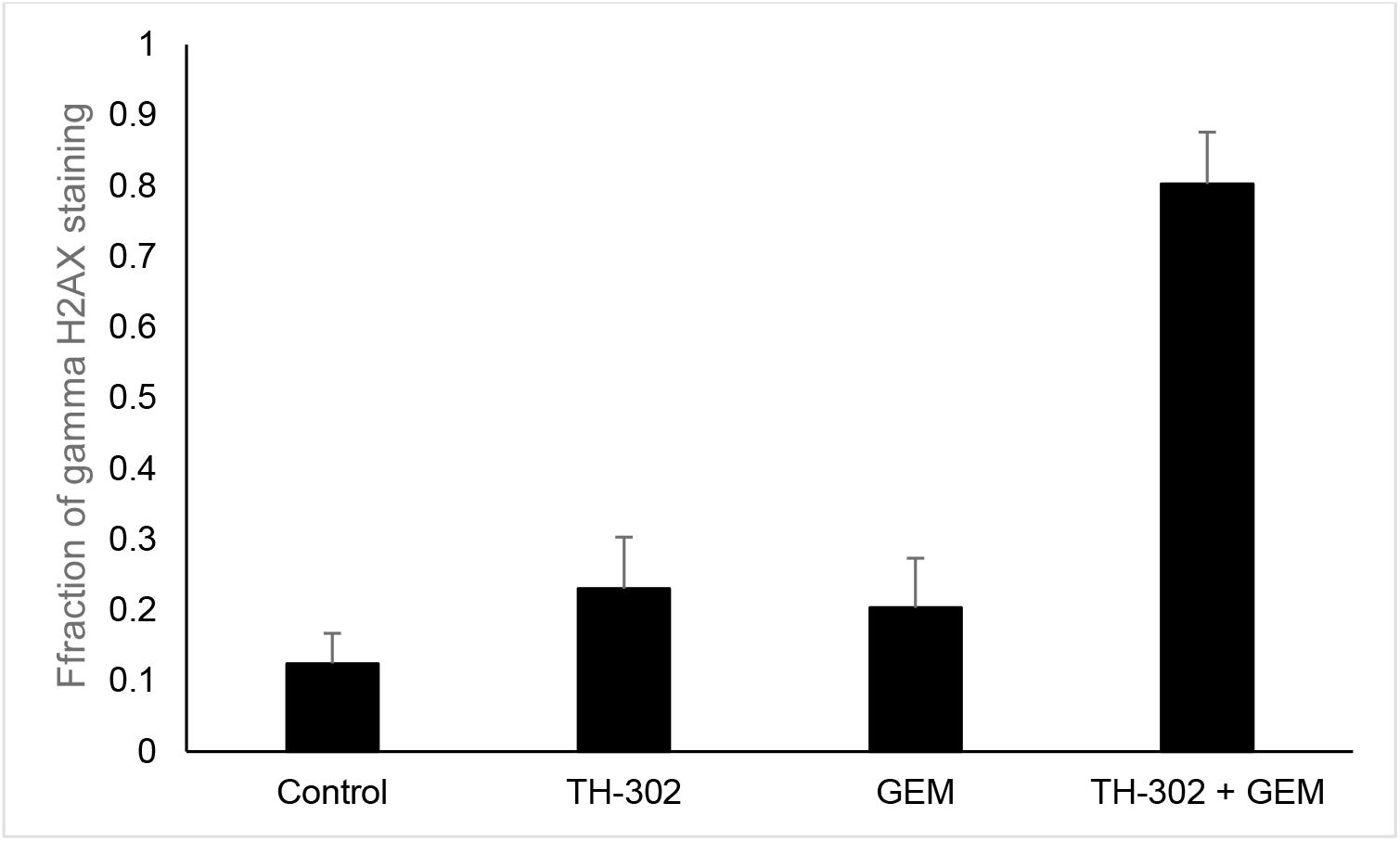
Fraction of adherent SU 86.86 cells showing double stranded DNA breaks by γH2AX staining. Cells were treated for 24 hours with sublethal concentrations of GEM and/or evofosfamide (IC20, GEM=2000 nM and evofosfamide=100 μM) and stained overnight at 4° C with anti-γH2AX antibody

## Discussion

Evofosfamide and GEM are known to act synergistically in PDAC xenografts, [21, 24] but the underlying mechanism remains unknown. Our data showed that both MIA Paca-2 and SU.86.86 cells had a similar sensitivity to Evo and MIA Paca-2 cells had a lower IC_50_ values with GEM than SU.86.86 cells. On the other hand, SU.86.86 tumor was not sensitive to evofosfamide and MIA Paca-2 tumor was resistant to GEM and SU.86.86 tumor was resistant to evofosfamide in in vivo assessment. This discrepancy is assumed to be the result of different physiologic profiles of TME. Similar to the observation, the effect of combination treatment *in vivo* was not consistent with in vitro cytotoxicity assay results. Although the combination treatment by evofosfamide and GEM showed synergistical effect only in SU.86.86 cells, such effect was observed in both type tumors *in vivo*.

The two tumors have different levels of tumor oxygenation [38] and extracellular matrix proteins [39]. MIA Paca-2 tumor is poorly differentiated tumor with low tumor oxygenation, while SU.86.86 tumor is moderately differentiated tumor with relatively higher tumor oxygenation due to its better constructed vasculature. The tumor oxygenation profile is associated with the drug sensitivity of the tumors. Evofosfamide exerts its cytotoxicity under hypoxic state [22, 27, 28, 38] and GEM is, in general, more potent in well perfused tissue [26, 40].

While both are PDAC tumors, MIA Paca-2 and SU86.86 xenografts differ strongly in the tumor microenvironment. MIA Paca-2 tumors are poorly differentiated with immature, atonal vasculature that contributes to hypoxic environment, while SU.86.86 xenografts are moderately differentiated with relatively higher tumor oxygenation due to their better constructed vasculature.[38, 40, 41] The tumor oxygenation profile is associated with the relative drug sensitivity of the tumors. Evofosfamide exerts its cytotoxicity under a hypoxic state [22, 27, 28] and GEM is, in general, more potent in well perfused tissue.[42] While these two mechanisms appear to be acting in opposing directions, there is an apparent bilateral action of each drug on the other – even when GEM (MIA Paca-2) or evofosfamide (SU86.86) is ineffective the combination therapy is more effective than the additive effect of both drugs (Figure 1).

The origin of this synergy seems to differ in the two cell lines. In MIAPaCa-2 xenografts, a clear mechanistic rationale for the effect of GEM on evofosfamide is provided by the increase in the hypoxic fraction of the tumor after GEM, although the physiological origin of this increase is elusive. As with other nucleoside analogues,[43] there are a number of vascular effects associated with gemcitabine treatment, primarily peripheral edema from extravasation of plasma,[44] possibly due to gemcitabine’s toxic effect on the endothelial cells lining capillaries.[45, 46] Gemcitabine induced capillary hyperpermeability is broadly consistent with the hemodynamic features observed in Figure 4 – blood volume remains constant after treatment while oxygen levels in the hypoxic interior of the tumor decrease but the periphery remains largely unchanged. In the flow limited situation expected to prevail in the constricted and hypovascularized PDAC tumor microenvironment, oxygen delivery in leaky vasculature is expected to be reduced as oxygen leaks into the extravascular space before reaching the venous side of the capillary bed.[33] Although speculative, this mechanism has some support from study using Overhauser MRI which showed high vascular permeability is sufficient to cause hypoxia in SCCVII tumors regardless of the degree of blood perfusion.[47]

While an increase in the activity of evofosfamide due to a rise in hypoxia through the action of GEM is plausible for MIA PaCa-2, this mechanism is less likely for SU86.86. In contrast to MIA Paca-2, most of the synergy between evofosfamide and GEM was present in cell culture, which would not be expected if a change in hypoxia levels was the cause. In cell culture, the synergy is roughly reciprocal – the additive effect of a sublethal dose of GEM in the presence of a near-lethal dose of Evofosfamide is roughly equivalent to the additive effect of a sublethal dose of Evofosfamide in the presence of a near-lethal dose of GEM (Figure 2F). This symmetry suggests a bilateral action and the possibility a shared pathway may be active in both. The double stranded DNA lesions caused by evofosfamide are almost exclusively repaired by homologous recombination, as mutations in key proteins in other pathways have almost no effect on evofosfamide activity.[28] Gemcitabine inhibits DNA repair in a number of ways.[34] Most importantly for evofosfamide, gemcitabine inhibits homologous recombination by stalling replication forks, either thorough chain termination when GEM is incorporated into replicating DNA[35] or by suppression of the activity of the RAD51 protein that controls strand pairing during homologous recombination.[36] This creates a situation of replication stress and arrest in the G2/M cell cycle.[48, 49] In this respect, evofosfamide may not be dissimilar to cisplatin, another DNA cross-linking agent commonly used in combination with GEM with a documented synergy that stems from inhibition of homologous recombination and nucleotide excision repair by GEM.[50] Evofosfamide also increased blood volume within the tumor, which may be an indication of improved transport of GEM in SU86.86. [51]

## Author Contributions

**Yasunori Otowa:** Investigation, Methodology, Writing – original draft, **Shun Kishimoto**: Investigation, Conceptualization, Methodology, Supervision, Writing – original draft. **Yu Saida:** Investigation, Methodology. **Kota Yamashita:** Investigation, **Kazutoshi Yamamoto:** Investigation. **Gadisetti VR Chandramouli:** Formal analysis, Investigation, Software. **Nallathamby Devasahayam:** Resources, Investigation. **James B. Mitchell:** Project administration, Supervision, Funding acquisition. **Murali C. Krishna:** Conceptualization, Project administration, Methodology, Supervision, Funding acquisition. **Jeffery R. Brender:** Conceptualization, Methodology, Formal analysis, Writing – original draft Writing – review & editing.

## Figure Legends

**Figure S1.**
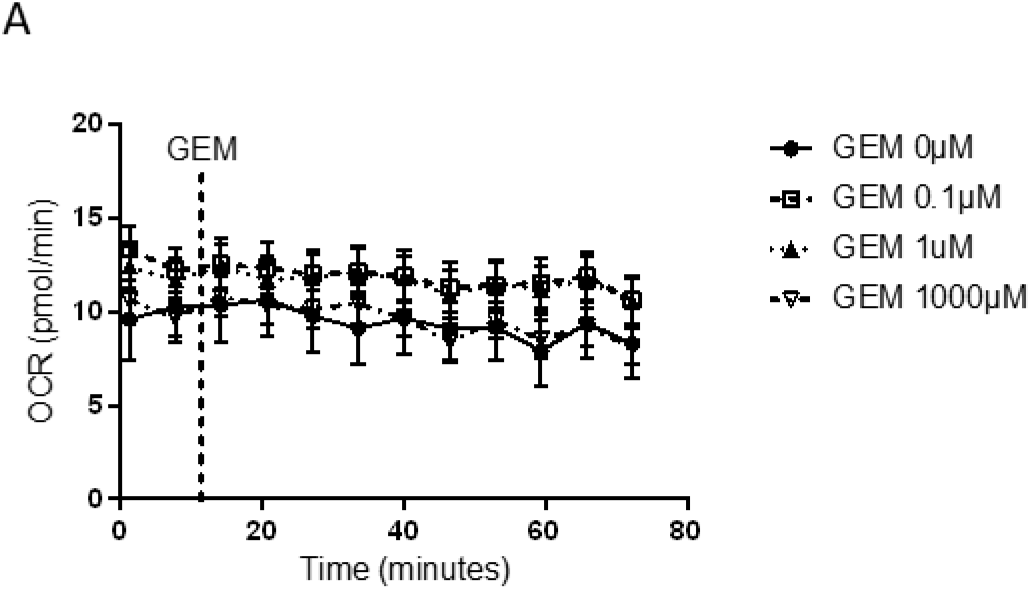
Oxygenation consumption does not change after GEM treatment in MIA Paca-2 tumors. **(A)** Oxygenation consumption rate was not affected by treatment of GEM. **(B)** Extracellular acidification rate (ECAR) was decreased after treated with GEM and recovered when cells were treated with low concentration of GEM. On the other hand, ECAR remained low when cells were treated with high concentration of GEM. Data are shown as mean ± SE.

**Figure S2.**
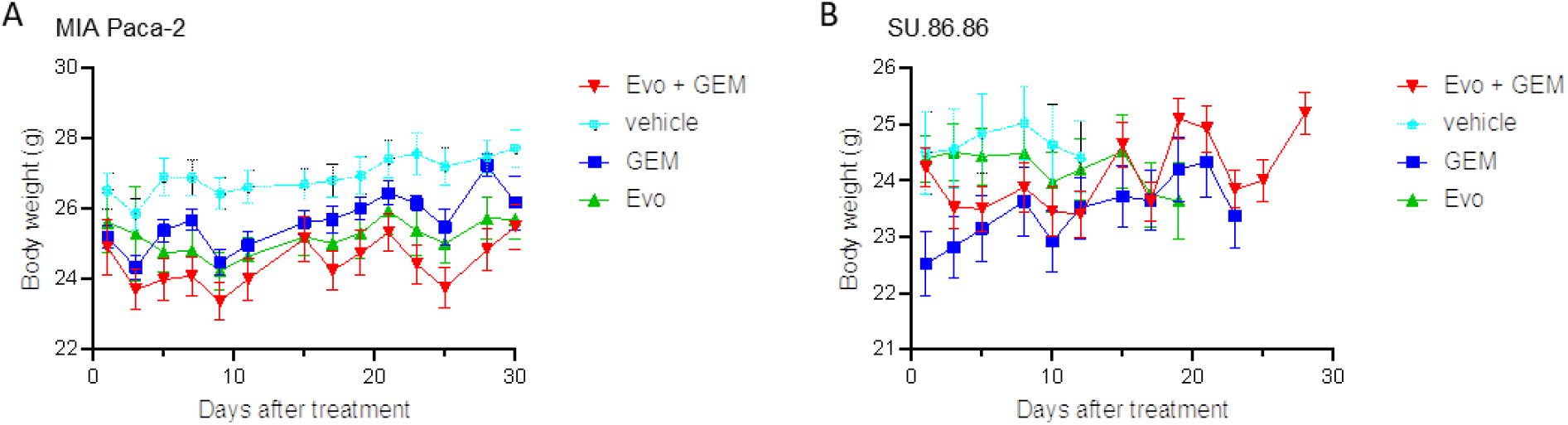
Body weight does not change after treatment. The body weight was similar between treatment groups in MIA Paca-2 tumors **(A)** and SU.86.86 tumors **(B)**. Data are shown as mean ± SE.

**Figure S3.**
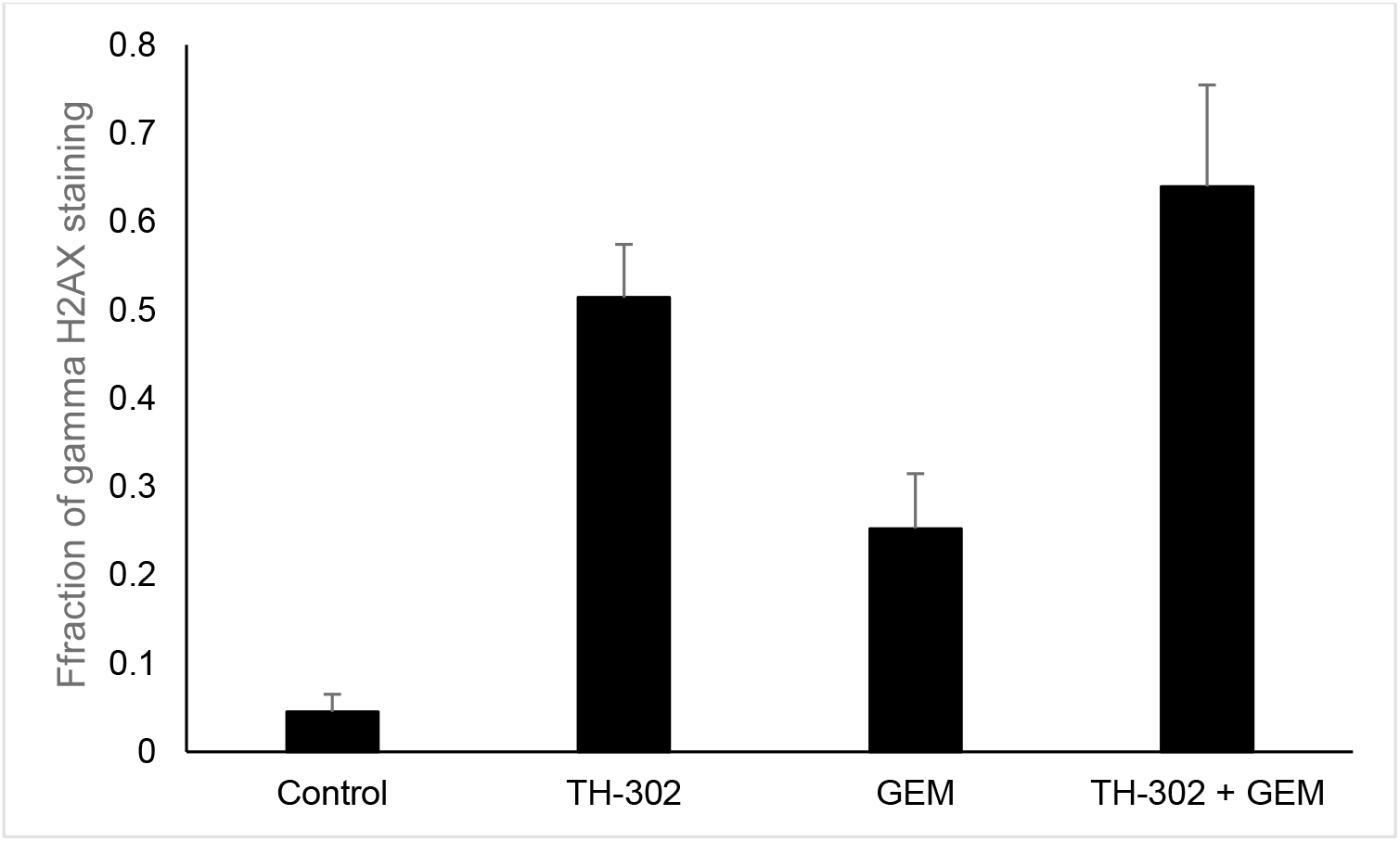
Fraction of adherent MIA PaCa-2 cells showing double stranded DNA breaks by γH2AX staining. Cells were treated for 24 hours with sublethal concentrations of GEM and/or evofosfamide (GEM=50 nM and evofosfamide=100 μM) and stained overnight at 4° C with anti-γH2AX antibody.

## References

[1] G.B.D.P.C. Collaborators, The global, regional, and national burden of pancreatic cancer and its attributable risk factors in 195 countries and territories, 1990-2017: a systematic analysis for the Global Burden of Disease Study 2017, Lancet Gastroenterol Hepatol, 4 (2019) 934–947.

[2] F. Giovinazzo, G. Turri, S. Zanini, G. Butturini, A. Scarpa, C. Bassi, Clinical implications of biological markers in Pancreatic Ductal Adenocarcinoma, Surg Oncol, 21 (2012) e171–182.

[3] H.A. Burris, 3rd, M.J. Moore, J. Andersen, M.R. Green, M.L. Rothenberg, M.R. Modiano, M.C. Cripps, R.K. Portenoy, A.M. Storniolo, P. Tarassoff, R. Nelson, F.A. Dorr, C.D. Stephens, D.D. Von Hoff, Improvements in survival and clinical benefit with gemcitabine as first-line therapy for patients with advanced pancreas cancer: a randomized trial, J Clin Oncol, 15 (1997) 2403–2413.

[4] M.P. Kim, G.E. Gallick, Gemcitabine resistance in pancreatic cancer: picking the key players, Clin Cancer Res, 14 (2008) 1284–1285.

[5] T. Brodowicz, R.M. Wolfram, W.J. Kostler, S. Tomek, I. Vaclavik, G.G. Steger, B. Teleky, R. Fugger, R. Jakesz, C.C. Zielinski, Phase II study of gemcitabine in combination with cisplatin in patients with locally advanced and/or metastatic pancreatic cancer, Anticancer Drugs, 11 (2000) 623–628.

[6] H.Q. Xiong, A. Rosenberg, A. LoBuglio, W. Schmidt, R.A. Wolff, J. Deutsch, M. Needle, J.L. Abbruzzese, Cetuximab, a monoclonal antibody targeting the epidermal growth factor receptor, in combination with gemcitabine for advanced pancreatic cancer: a multicenter phase II Trial, J Clin Oncol, 22 (2004) 2610–2616.

[7] C. Louvet, R. Labianca, P. Hammel, G. Lledo, M.G. Zampino, T. Andre, A. Zaniboni, M. Ducreux, E. Aitini, J. Taieb, R. Faroux, C. Lepere, A. de Gramont, Gercor, Giscad, Gemcitabine in combination with oxaliplatin compared with gemcitabine alone in locally advanced or metastatic pancreatic cancer: results of a GERCOR and GISCAD phase III trial, J Clin Oncol, 23 (2005) 3509–3516.

[8] V. Heinemann, D. Quietzsch, F. Gieseler, M. Gonnermann, H. Schonekas, A. Rost, H. Neuhaus, C. Haag, M. Clemens, B. Heinrich, U. Vehling-Kaiser, M. Fuchs, D. Fleckenstein, W. Gesierich, D. Uthgenannt, H. Einsele, A. Holstege, A. Hinke, A. Schalhorn, R. Wilkowski, Randomized phase III trial of gemcitabine plus cisplatin compared with gemcitabine alone in advanced pancreatic cancer, J Clin Oncol, 24 (2006) 3946–3952.

[9] T. Okusaka, H. Ishii, A. Funakoshi, H. Ueno, J. Furuse, T. Sumii, A phase I/II study of combination chemotherapy with gemcitabine and 5-fluorouracil for advanced pancreatic cancer, Jpn J Clin Oncol, 36 (2006) 557–563.

[10] T. Conroy, F. Desseigne, M. Ychou, O. Bouche, R. Guimbaud, Y. Becouarn, A. Adenis, J.L. Raoul, S. Gourgou-Bourgade, C. de la Fouchardiere, J. Bennouna, J.B. Bachet, F. Khemissa-Akouz, D. Pere-Verge, C. Delbaldo, E. Assenat, B. Chauffert, P. Michel, C. Montoto-Grillot, M. Ducreux, U. Groupe Tumeurs Digestives of, P. Intergroup, FOLFIRINOX versus gemcitabine for metastatic pancreatic cancer, N Engl J Med, 364 (2011) 1817–1825.

[11] P. Vaupel, Tumor microenvironmental physiology and its implications for radiation oncology, Semin Radiat Oncol, 14 (2004) 198–206.

[12] T. Samkharadze, M. Erkan, C. Reiser-Erkan, I.E. Demir, B. Kong, G.O. Ceyhan, C.W. Michalski, I. Esposito, H. Friess, J. Kleeff, Pigment epithelium-derived factor associates with neuropathy and fibrosis in pancreatic cancer, Am J Gastroenterol, 106 (2011) 968–980.

[13] M. Erkan, C. Reiser-Erkan, C.W. Michalski, S. Deucker, D. Sauliunaite, S. Streit, I. Esposito, H. Friess, J. Kleeff, Cancer-stellate cell interactions perpetuate the hypoxia-fibrosis cycle in pancreatic ductal adenocarcinoma, Neoplasia, 11 (2009) 497–508.

[14] O. Kisker, S. Onizuka, J. Banyard, T. Komiyama, C.M. Becker, E.G. Achilles, C.M. Barnes, M.S. O’Reilly, J. Folkman, S.R. Pirie-Shepherd, Generation of multiple angiogenesis inhibitors by human pancreatic cancer, Cancer Res, 61 (2001) 7298–7304.

[15] C.A. O’Mahony, D. Albo, G.P. Tuszynski, D.H. Berger, Transforming growth factor-beta 1 inhibits generation of angiostatin by human pancreatic cancer cells, Surgery, 124 (1998) 388–393.

[16] Q. Chang, I. Jurisica, T. Do, D.W. Hedley, Hypoxia predicts aggressive growth and spontaneous metastasis formation from orthotopically grown primary xenografts of human pancreatic cancer, Cancer Res, 71 (2011) 3110–3120.

[17] J. Brender, Y. Saida, N. Devasahayam, M.K. Cherukuri, S. Kishimoto, Hypoxia Imaging as a Guide for Hypoxia Modulated and Hypoxia Activated Therapy, Antioxid Redox Signal, (2021).

[18] Y. Takakusagi, S. Kishimoto, S. Naz, S. Matsumoto, K. Saito, C.P. Hart, J.B. Mitchell, M.C. Krishna, Radiotherapy Synergizes with the Hypoxia-Activated Prodrug Evofosfamide: In Vitro and In Vivo Studies, Antioxid Redox Signal, 28 (2018) 131–140.

[19] E. Van Cutsem, H.J. Lenz, J. Furuse, J. Tabernero, V. Heinemann, T. Ioka, I. Bazin, M. Ueno, T. Csoszi, H. Wasan, B. Melichar, P. Karasek, T.M. Macarulla, C. Guillen, E. Kalinka-Warzocha, Z. Horvath, H. Prenen, M. Schlichting, A. Ibrahim, J.C. Bendell, MAESTRO: A randomized, double-blind phase III study of evofosfamide (Evo) in combination with gemcitabine (Gem) in previously untreated patients (pts) with metastatic or locally advanced unresectable pancreatic ductal adenocarcinoma (PDAC), Journal of Clinical Oncology, 34 (2016).

[20] C. Hajj, J. Russell, C.P. Hart, K.A. Goodman, M.A. Lowery, A. Haimovitz-Friedman, J.O. Deasy, J.L. Humm, A Combination of Radiation and the Hypoxia-Activated Prodrug Evofosfamide (TH-302) is Efficacious against a Human Orthotopic Pancreatic Tumor Model, Transl Oncol, 10 (2017) 760–765.

[21] Q. Liu, J.D. Sun, J. Wang, D. Ahluwalia, A.F. Baker, L.D. Cranmer, D. Ferraro, Y. Wang, J.X. Duan, W.S. Ammons, J.G. Curd, M.D. Matteucci, C.P. Hart, TH-302, a hypoxia-activated prodrug with broad in vivo preclinical combination therapy efficacy: optimization of dosing regimens and schedules, Cancer Chemother Pharmacol, 69 (2012) 1487–1498.

[22] S. Matsumoto, S. Kishimoto, K. Saito, Y. Takakusagi, J.P. Munasinghe, N. Devasahayam, C.P. Hart, R.J. Gillies, J.B. Mitchell, M.C. Krishna, Metabolic and Physiologic Imaging Biomarkers of the Tumor Microenvironment Predict Treatment Outcome with Radiation or a Hypoxia-Activated Prodrug in Mice, Cancer Res, 78 (2018) 3783–3792.

[23] S.G. Peeters, C.M. Zegers, R. Biemans, N.G. Lieuwes, R.G. van Stiphout, A. Yaromina, J.D. Sun, C.P. Hart, A.D. Windhorst, W. van Elmpt, L.J. Dubois, P. Lambin, TH-302 in Combination with Radiotherapy Enhances the Therapeutic Outcome and Is Associated with Pretreatment [18F]HX4 Hypoxia PET Imaging, Clin Cancer Res, 21 (2015) 2984–2992.

[24] J.D. Sun, Q. Liu, D. Ahluwalia, W. Li, F. Meng, Y. Wang, D. Bhupathi, A.S. Ruprell, C.P. Hart, Efficacy and safety of the hypoxia-activated prodrug TH-302 in combination with gemcitabine and nab-paclitaxel in human tumor xenograft models of pancreatic cancer, Cancer Biol Ther, 16 (2015) 438–449.

[25] Y. Takakusagi, S. Matsumoto, K. Saito, M. Matsuo, S. Kishimoto, J.W. Wojtkowiak, W. DeGraff, A.H. Kesarwala, R. Choudhuri, N. Devasahayam, S. Subramanian, J.P. Munasinghe, R.J. Gillies, J.B. Mitchell, C.P. Hart, M.C. Krishna, Pyruvate induces transient tumor hypoxia by enhancing mitochondrial oxygen consumption and potentiates the anti-tumor effect of a hypoxia-activated prodrug TH-302, PLoS One, 9 (2014) e107995.

[26] J.W. Wojtkowiak, H.C. Cornnell, S. Matsumoto, K. Saito, Y. Takakusagi, P. Dutta, M. Kim, X. Zhang, R. Leos, K.M. Bailey, G. Martinez, M.C. Lloyd, C. Weber, J.B. Mitchell, R.M. Lynch, A.F. Baker, R.A. Gatenby, K.A. Rejniak, C. Hart, M.C. Krishna, R.J. Gillies, Pyruvate sensitizes pancreatic tumors to hypoxia-activated prodrug TH-302, Cancer Metab, 3 (2015) 2.

[27] S. Kishimoto, J.R. Brender, G.V.R. Chandramouli, Y. Saida, K. Yamamoto, J.B. Mitchell, M.C. Krishna, Hypoxia-Activated Prodrug Evofosfamide Treatment in Pancreatic Ductal Adenocarcinoma Xenografts Alters the Tumor Redox Status to Potentiate Radiotherapy, Antioxid Redox Signal, (2020).

[28] F. Meng, J.W. Evans, D. Bhupathi, M. Banica, L. Lan, G. Lorente, J.X. Duan, X. Cai, A.M. Mowday, C.P. Guise, A. Maroz, R.F. Anderson, A.V. Patterson, G.C. Stachelek, P.M. Glazer, M.D. Matteucci, C.P. Hart, Molecular and cellular pharmacology of the hypoxia-activated prodrug TH-302, Mol Cancer Ther, 11 (2012) 740–751.

[29] T. Arumugam, V. Ramachandran, K.F. Fournier, H. Wang, L. Marquis, J.L. Abbruzzese, G.E. Gallick, C.D. Logsdon, D.J. McConkey, W. Choi, Epithelial to mesenchymal transition contributes to drug resistance in pancreatic cancer, Cancer Res, 69 (2009) 5820–5828.

[30] S. Matsumoto, S. Batra, K. Saito, H. Yasui, R. Choudhuri, C. Gadisetti, S. Subramanian, N. Devasahayam, J.P. Munasinghe, J.B. Mitchell, M.C. Krishna, Antiangiogenic agent sunitinib transiently increases tumor oxygenation and suppresses cycling hypoxia, Cancer Res, 71 (2011) 6350–6359.

[31] S. Matsumoto, F. Hyodo, S. Subramanian, N. Devasahayam, J. Munasinghe, E. Hyodo, C. Gadisetti, J.A. Cook, J.B. Mitchell, M.C. Krishna, Low-field paramagnetic resonance imaging of tumor oxygenation and glycolytic activity in mice, J Clin Invest, 118 (2008) 1965–1973.

[32] S. Matsumoto, K. Saito, H. Yasui, H.D. Morris, J.P. Munasinghe, M. Lizak, H. Merkle, J.H. Ardenkjaer-Larsen, R. Choudhuri, N. Devasahayam, S. Subramanian, A.P. Koretsky, J.B. Mitchell, M.C. Krishna, EPR oxygen imaging and hyperpolarized 13C MRI of pyruvate metabolism as noninvasive biomarkers of tumor treatment response to a glycolysis inhibitor 3-bromopyruvate, Magn Reson Med, 69 (2013) 1443–1450.

[33] S. Matsumoto, K. Saito, Y. Takakusagi, M. Matsuo, J.P. Munasinghe, H.D. Morris, M.J. Lizak, H. Merkle, K. Yasukawa, N. Devasahayam, S. Suburamanian, J.B. Mitchell, M.C. Krishna, In vivo imaging of tumor physiological, metabolic, and redox changes in response to the anti-angiogenic agent sunitinib: longitudinal assessment to identify transient vascular renormalization, Antioxid Redox Signal, 21 (2014) 1145–1155.

[34] S. Muller, DNA Damage-inducing Compounds: Unraveling their Pleiotropic Effects Using High Throughput Sequencing, Curr Med Chem, 24 (2017) 1558–1585.

[35] W. Plunkett, P. Huang, V. Gandhi, Preclinical characteristics of gemcitabine, Anticancer Drugs, 6 Suppl 6 (1995) 7–13.

[36] S. Kobashigawa, K. Morikawa, H. Mori, G. Kashino, Gemcitabine Induces Radiosensitization Through Inhibition of RAD51-dependent Repair for DNA Double-strand Breaks, Anticancer Res, 35 (2015) 2731–2737.

[37] L.J. Mah, A. El-Osta, T.C. Karagiannis, gammaH2AX: a sensitive molecular marker of DNA damage and repair, Leukemia, 24 (2010) 679–686.

[38] K. Yamamoto, J.R. Brender, T. Seki, S. Kishimoto, N. Oshima, R. Choudhuri, S.S. Adler, E.M. Jagoda, K. Saito, N. Devasahayam, P.L. Choyke, J.B. Mitchell, M.C. Krishna, Molecular Imaging of the Tumor Microenvironment Reveals the Relationship between Tumor Oxygenation, Glucose Uptake, and Glycolysis in Pancreatic Ductal Adenocarcinoma, Cancer Res, 80 (2020) 2087–2093.

[39] M. Lohr, B. Trautmann, M. Gottler, S. Peters, I. Zauner, B. Maillet, G. Kloppel, Human ductal adenocarcinomas of the pancreas express extracellular matrix proteins, Br J Cancer, 69 (1994) 144–151.

[40] K.M. Bailey, H.H. Cornnell, A. Ibrahim-Hashim, J.W. Wojtkowiak, C.P. Hart, X. Zhang, R. Leos, G.V. Martinez, A.F. Baker, R.J. Gillies, Evaluation of the “steal” phenomenon on the efficacy of hypoxia activated prodrug TH-302 in pancreatic cancer, PLoS One, 9 (2014) e113586.

[41] R. Gradiz, H.C. Silva, L. Carvalho, M.F. Botelho, A. Mota-Pinto, MIA PaCa-2 and PANC-1 - pancreas ductal adenocarcinoma cell lines with neuroendocrine differentiation and somatostatin receptors, Sci Rep, 6 (2016) 21648.

[42] W. Tang, W. Liu, H.M. Li, Q.F. Wang, C.X. Fu, X.H. Wang, L.P. Zhou, W.J. Peng, Quantitative dynamic contrast-enhanced MR imaging for the preliminary prediction of the response to gemcitabine-based chemotherapy in advanced pancreatic ductal carcinoma, Eur J Radiol, 121 (2019) 108734.

[43] J. Herrmann, E.H. Yang, C.A. Iliescu, M. Cilingiroglu, K. Charitakis, A. Hakeem, K. Toutouzas, M.A. Leesar, C.L. Grines, K. Marmagkiolis, Vascular Toxicities of Cancer Therapies The Old and the New - An Evolving Avenue, Circulation, 133 (2016) 1272–1289.

[44] C.A. Dasanu, Gemcitabine: vascular toxicity and prothrombotic potential, Expert Opin Drug Saf, 7 (2008) 703–716.

[45] W. Wu, Q. Xia, R.J. Luo, Z.Q. Lin, P. Xue, In vitro Study of the Antagonistic Effect of Low-dose Liquiritigenin on Gemcitabine-induced Capillary Leak Syndrome in Pancreatic Adenocarcinoma via Inhibiting ROS-Mediated Signalling Pathways, Asian Pac J Cancer Prev, 16 (2015) 4369–4376.

[46] A.J. van Hell, A. Haimovitz-Friedman, Z. Fuks, W.D. Tap, R. Kolesnick, Gemcitabine kills proliferating endothelial cells exclusively via acid sphingomyelinase activation, Cell Signal, 34 (2017) 86–91.

[47] S. Matsumoto, H. Yasui, S. Batra, Y. Kinoshita, M. Bernardo, J.P. Munasinghe, H. Utsumi, R. Choudhuri, N. Devasahayam, S. Subramanian, J.B. Mitchell, M.C. Krishna, Simultaneous imaging of tumor oxygenation and microvascular permeability using Overhauser enhanced MRI, P Natl Acad Sci USA, 106 (2009) 17898–17903.

[48] Y. Jiang, H. Dai, Y. Li, J. Yin, S. Guo, S.Y. Lin, D.J. McGrail, PARP inhibitors synergize with gemcitabine by potentiating DNA damage in non-small-cell lung cancer, Int J Cancer, 144 (2019) 1092–1103.

[49] H. Gaillard, T. Garcia-Muse, A. Aguilera, Replication stress and cancer, Nat Rev Cancer, 15 (2015) 276–289.

[50] L. Pellegrini, D.S. Yu, T. Lo, S. Anand, M. Lee, T.L. Blundell, A.R. Venkitaraman, Insights into DNA recombination from the structure of a RAD51-BRCA2 complex, Nature, 420 (2002) 287–293.

[51] T. Stylianopoulos, R.K. Jain, Combining two strategies to improve perfusion and drug delivery in solid tumors, Proc Natl Acad Sci U S A, 110 (2013) 18632–18637.

